# Distinct sex differences in the production of steroids and neuropeptides in the adult zebrafish brain-pituitary gonadal axis

**DOI:** 10.1101/2025.06.22.660951

**Authors:** Di Peng, Chunyu Lu, Vance L. Trudeau

## Abstract

Zebrafish are increasingly used as experimental models in studies of human disease, environmental toxicology, and reproductive biology. However, sex differences in hormone production are rarely examined, despite evidence from gene mutation studies indicating differential effects in females and males. The emerging reproductive peptide secretoneurin (SN) has not been quantitatively compared between sexes in any species. Here, we employed a newly developed extraction and LC-MS method to simultaneously quantify and compare levels of five steroids and thirteen peptides in brain, pituitary, and gonads. As expected, testosterone (T) and 11-ketotestosterone (11-KT) were higher in male tissues, while estrone (E1) and estradiol (E2) were elevated in the female pituitary/ovary and brain, respectively. Estriol (E3) was more abundant in testes. Gonadotropin-releasing hormones Gnrh2 and Gnrh3 were notably higher in testes. Oxytocin (isotocin) and vasopressin (vasotocin) were elevated in the female brain and in testes. Kisspeptins 1 and 2 also showed higher levels in testes. Similarly, SNa and SNb were more abundant in the female brain and pituitary, and markedly higher in testes than ovaries. Several smaller SN fragments were detected at low levels, with patterns suggesting sex-specific enzymatic processing. These findings reveal pronounced sex differences in both classical and emerging reproductive hormones and identify the SN peptide family as a new component of the brain–pituitary–gonadal axis. This dataset provides a foundation for future studies on sexually differentiated neuropeptide production and function across tissues.

## Introduction

The coordinated release of hormones from the brain–pituitary–gonadal (BPG) axis regulates ovarian and testicular functions in females and males, thus determining fertility in vertebrates. Reproductive hormones include the classical pituitary glycoproteins—luteinising hormone (Lh) and follicle-stimulating hormone (Fsh)—and gonadal steroids, as well as an array of hypothalamic neuropeptides and neurotransmitters implicated in controlling pituitary hormone release (Trudeau 2022). In mammals, hypothalamic gonadotropin-releasing hormone (Gnrh1) neurons project to the highly vascularised median eminence, where this decapeptide is released in sex-specific patterns into the portal circulation to regulate pituitary functions. For instance, there is an Lh surge triggering ovulation in females (Goodman et al., 2022), and pulsatile Lh release with parallel testosterone production in males (Sanford et al., 1984). Gonadal steroids in turn exert feedback at the hypothalamic and pituitary levels to finely regulate Lh production in both sexes. Other neuropeptides, such as kisspeptin and dynorphin, may modulate Gnrh release or have additional roles (Moore et al., 2023). In teleost fish, numerous neuropeptides also regulate pituitary function. However, unlike mammals, teleosts lost the median eminence over evolutionary time. Consequently, both females and males exhibit extensive direct innervation of the anterior pituitary, in addition to classical innervation of the posterior pituitary (Trudeau & Somoza, 2020; Muñoz-Cueto et al., 2020).

Overt sexual dimorphism refers to the presence of two distinct forms in the sexes, whereas sex differences refer to endpoints existing on a continuum, with differing averages between males and females (McCarthy et al., 2012). Significant sex differences exist in both reproductive and non-reproductive physiology, often reflected in hormone production patterns. The influence of sex is far-reaching, yet historically, females have been underrepresented in research. Over a decade ago, several funding agencies mandated the inclusion of sex as a variable in biomedical studies (McCarthy et al., 2012). Hormonal and genetic sex differences have been most extensively studied in mammals (Marrocco & McEwen, 2016; Jennings & de Lecea, 2020) and birds (Adkins-Regan, 2007).

In teleosts, sex differences in endocrine systems have been studied in species such as goldfish (Bosma et al., 2001; Qi, Zhou et al., 2013) and medaka (Ohya & Hayashi, 2006; Kawabata et al., 2012; Yamashita et al., 2021). Recent work also identified sex differences in Gnrh3 neuron innervation by tachykinin1 neurons in zebrafish (Ogawa et al., 2020). Emerging studies are revealing sex differences in gene expression in fish gonads and brains (Dai et al., 2023; Rajendiran et al., 2021). These investigations typically focus on mechanisms of gonadal or brain sex differentiation. A few have compared female and male pituitaries (e.g., Margolis-Kazan & Schreibman, 1984; Mun et al., 2022; Rao et al., 1996; Royan et al., 2021; Yamashita et al., 2021). However, studies explicitly comparing hormone production in the brain, pituitary, and gonads of adult post-pubertal female and male teleosts are scarce (Zhai et al., 2022).

This scarcity is surprising for several reasons. Many fish species are of high socio-economic value as food, and understanding sex differences is key to improving aquaculture— for instance, by identifying endocrine mechanisms causing one sex to grow faster. Teleosts represent the largest vertebrate group and exhibit exceptional diversity in reproductive strategies (Trudeau & Somoza, 2024). They are excellent models for exploring the significance of genome duplications and the roles of paralogous genes in reproductive physiology and evolutionary success (Dufour et al., 2020). Small gonochoristic species such as zebrafish (*Danio rerio*) are also widely used models in studies of human disease, environmental science, and reproductive biology. Yet we know surprisingly little about sex-specific differences in hormone levels.

To address this gap, we investigated neuropeptide and gonadal steroid production in zebrafish. Using our newly developed simultaneous extraction and liquid chromatography– mass spectrometry (LC–MS) methods (Lu et al., 2023; Peng et al., 2025), we quantified five sex steroids and 13 neuropeptides in individual brain, pituitary, and gonad samples. Alongside classical hormones, we assessed two newer reproductive peptides, secretoneurin A (SNa) and secretoneurin B (SNb), which arise from selective prohormone convertase cleavage of secretogranin-2a (Scg2a) and Scg2b precursor proteins. These peptides are of interest because only 1 in 10 zebrafish pairs with double *scg2a/2b* frameshift mutations spawn (Mitchell, Zhang et al., 2020), and SNa injection robustly activates the HPG axis and induces ovulation (Peng et al., 2025).

## Material and Methods

### Experimental animal husbandry

Zebrafish (AB strain) were bred and kept in the University of Ottawa Aquatics core facility at the temperature of 28°C. The lights-on time was set for 14 hours from 0900h to 2300h. All animal experiments followed the guidelines of the Canadian Council on Animal Care for the use of animals in research, approved by the University of Ottawa Animal Care Protocol Review Committee. Sexually mature zebrafish at the age of 6-9 months post-fertilization (mpf) were selected and separated by sex at least two weeks to synchronize cycles, and to eliminate possible acute influences of one sex on the other through pheromones, and visual or tactile interactions. Females (n=12) and males (n=12) were dissections between 10:00-11:00. This is outside the normal nighttime periovulatory period to reduce possible variations due to gonadal cyclicity.

### Dissection and homogenization

The zebrafish were sacrificed by immersion in an ice bath and weighed. After spinal transection brain, pituitary, and gonads were dissected. Gonads were weighed for calculation of the gonadosomatic index (GSI; gonads weight/total body weight*100) while brains were weighed to calculate the brain-to-body mass ratio (brains weight/total body weight*100). The pituitaries were too small to weigh accurately so hormone levels are reported as total gland content. Dissected tissues were placed in 1.5 ml centrifuge tubes with homogenization buffer (90% MeOH, 9% water, and 1% acetic acid v/v/v) on ice and subjected to ultrasonic homogenization (Fisherbrand; Cat# FB120110) for 3-5 sec as previously reported (Lu et al. 2022). The homogenized samples were centrifuged at 4°C in 13,200 rpm for 10 min. After that, the supernatant was transferred into a new 1.5 ml tube and dehydrated in centrifugal vacuum concentrator (Labconco; Cat#7810014) at 45°C. Dehydrated samples were stored in -20°C until extraction.

### Solid phase extraction (SPE) and liquid chromatography-mass spectroscopy (LC-MS) analysis

The dehydrated supernatant samples were thawed from -20°C at room temperature and resuspended in loading buffer and subjected to SPE in a 96-well format. All details of the extraction, LS-MS methods, detection limits have been published (Lu et al. 2022). After collections the purified samples, the collection plate was transferred in the vacuum dried to be dehydrated and stored in -20°C till mass spectrometry detection. The LC-MS system is Nano flow LC-MS/MS analysis system driven by an Agilent 1100 capillary liquid chromatography system (running on micro mode) and a Thermo Fisher LTQ Velos Pro Orbitrap Elite mass spectrometer. The nano-HPLC was performed on 100 mm × 50 µm C18 columns with 75 µm ID loading column, and flow rates is set on 300 nL/min. LC elution is introduced into the mass analyzer by 2.5kV nano-electrospray ionization through a spray tip with a 1mm opening. Through the entire analysis, the Fourier Transform mass analyzer is set to collect m/z information between 150 m/z and 1,500 m/z on scan mode under mass resolution 60,000. The LC-MS result files were processed by Xcalibur software (Thermo Xcalibur 2.2 SP1. Build 48). Target compounds are identified by retention time (RSD≤2.5%), MS1 (10 ppm mass accuracy), and MS2 (0.5 m/z mass accuracy). Chromatograms were plotted based on the MS1 mass intensity with 20 ppm mass accuracy. Peak area was used to calculate the concentration of analytes in the original sample with an external standard curve. All peptides were prepared by standard Fmoc/tBu chemistry and purified by HPLC (>95%) in our laboratory (Lu et al., 2023) and steroids were purchased from Sigma-Aldrich Inc. (Table 1).

**Table 1.**
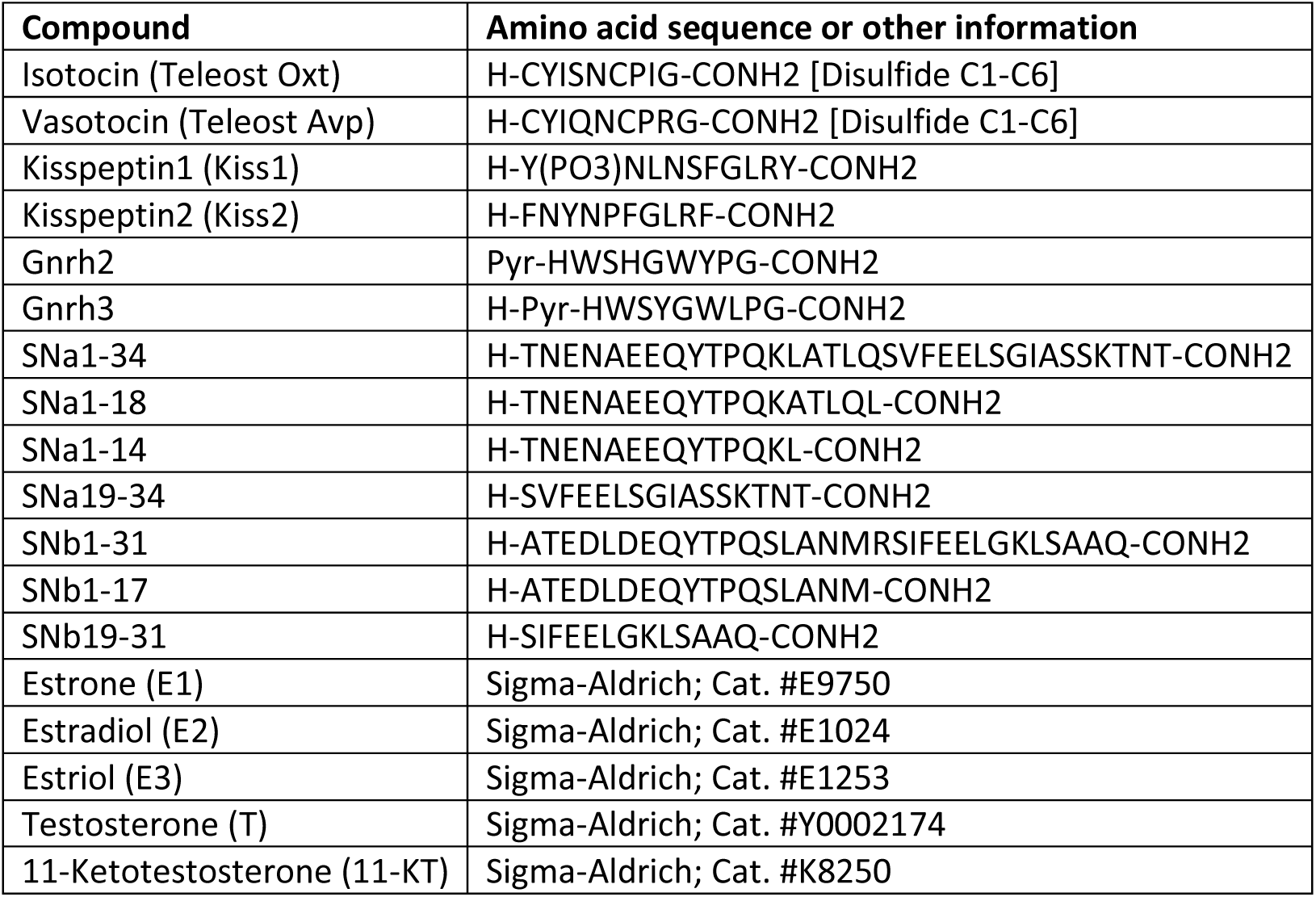
Standard compounds used for hormonal analysis by LC-MS.

### Gene and protein nomenclature

The zebrafish gene/protein nomenclature conventions (https://zfin.atlassian.net/wiki/spaces/general/pages/1818394635/ ZFIN+Zebrafish+Nomenclature+Conventions) have been used. The zebrafish nomenclature we have used for avp/Avp and oxt/Oxt is different from an earlier nomenclature of *avt*/Avt and *ist*/Ist (Mennigen et al., 2022). The reasoning for this is to highlight the homologous nature of *avp* and *oxt* gene families. It is implicitly understood that the teleost Avp and Oxt differ from mammalian and other vertebrate nonapeptide homologues in their amino residues (Table 1).

### Statistical analyses

The gonadosomatic index (gonad mass/body mass x 100) and brain-to-body mass index (brain mass/body mass X 100) were calculated and presented as mean ± SD. Levels of steroids and peptides are expressed as fmol/mg wet weight for brains and gonads and fmol/tissue for pituitaries as they are too small to weigh accurately. To assess the potential differences between the sexes in hormone metabolism, we calculated the product-to-parent hormone ratios in each tissue. For the sex steroids, these were ratios for T/11KT, T/E2, E1/E2, E1/E3 and E2/E3. We also calculated the ratios of smaller fragments relative to SNa1-34 (SNa-18, SNa1-14 and SNa19-34) and SNb1-31 (SNb1-17 and SNb19-31). The normality and log normality of the data were analyzed by Shapiro-Wilk test (GraphPad Prism 9 software). Consequently, sex differences in morphometric measures were analyzed using the two-tailed Student’s t-test. Differences in hormone levels were analysed using two-tailed Mann-Whitney U tests with calculated *z*-scores and *p*-values (https://www.socscistatistics.com/). Note that for graphs 2-8, data are plotted on a log scale for clarity since sex differences were large for some hormones.

For multivariate pattern exploration, Principal Component Analysis (PCA) was conducted using Python (v3.8+) and R (v4.1.0). Two datasets were prepared: one containing all quantified peptides and steroids, and another excluding SNa, SNb and their peptide fragments. PCA was performed using scikit-learn (v1.0.2) after data standardization, with analyses conducted for the complete dataset as well as stratified by sex and tissue type. PCA scatter plots (PC1 vs PC2) were visualized using ggplot2 (v3.3.5) in R, with samples distinguished by sex (circles for females, triangles for males) and explained variance ratios displayed on axis labels.

## Results

### Differences in sex steroid levels in female and male zebrafish

Female body mass was 623 ± 203 mg, being ∼1.4 times larger (*t*(11)= 3.01, *p*=0.006) than male body mass, which was 440 ± 58 mg. The GSI of females was 14.46 ± 3.34, while it was expectedly smaller in males and was 1.09 ± 0.15 (*t*(11)=13.47, *p*<0.001). In contrast, the brain-to-body mass index was 0.98 ± 0.30 in females, and somewhat larger in males, being 1.63 ± 0.18 (*t*(11)= -6.44, *p*<0.001).

Presented in Fig. 1 are the structures and interrelationships of the androgens and estrogens we quantified in the brains, pituitaries and gonads of both sexes. As can be seen in Fig. 2, there was a sex difference in testosterone (T) levels in the brains (z=-2.916, p=0.004) and gonads (*z*=-4.128, *p*<0.001) but not in the pituitary (*z*=-0.318, *p*=0.749). Based on the comparison of median values, brain T levels were 1.25 times higher in males than females. In the testes, T was 315.8 times higher than in the ovaries. Male-biased 11-KT levels were evident in all tissues (Fig. 2). The 11-KT levels were 7.8 (*z*=-3.782, *p*<0.001) and 14 (*z*=-4.128, *p*<0.001) times higher in the brain and pituitary of males compared to females. Similarly, testes levels of 11-KT were 10.8 times (*z*=-4.128, *p*<0.001) higher than in ovaries.

**Fig 1.**
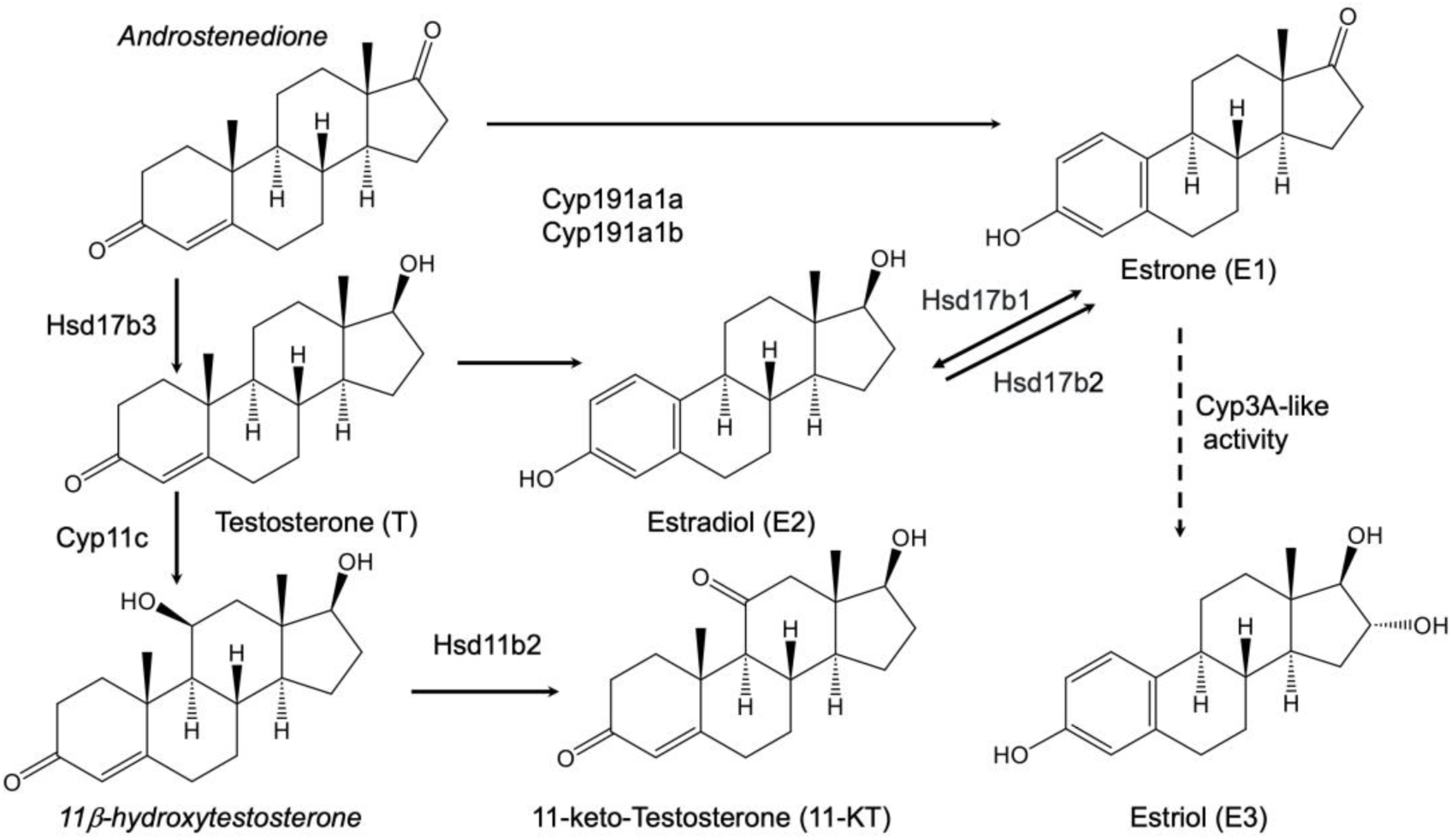
Simplified androgen and estrogen conversion pathways. Shown are the structures and interrelationships of the steroids measured in this study. Androstenedione and 11b-hydroxytestosterone are shown (in Italics) but were not measured. The main enzymes involved in steroid synthesis are hydroxysteroid 11-beta dehydrogenase type 2 (Hsd11b2), hydroxysteroid 17-beta dehydrogenase types 1, 2, 3 (Hsd17b1, Hsd17b2, Hsd17b3, cytochrome P450 (Cyp) family 11 type c1 (Cyp11c1), aromatase A (Cyp19a1a) and aromatase B (Cyp19a1b). Note that there are several possible routes (dotted line) for E3 production from E1 via a Cyp3A-like enzyme.

**Fig 2.**
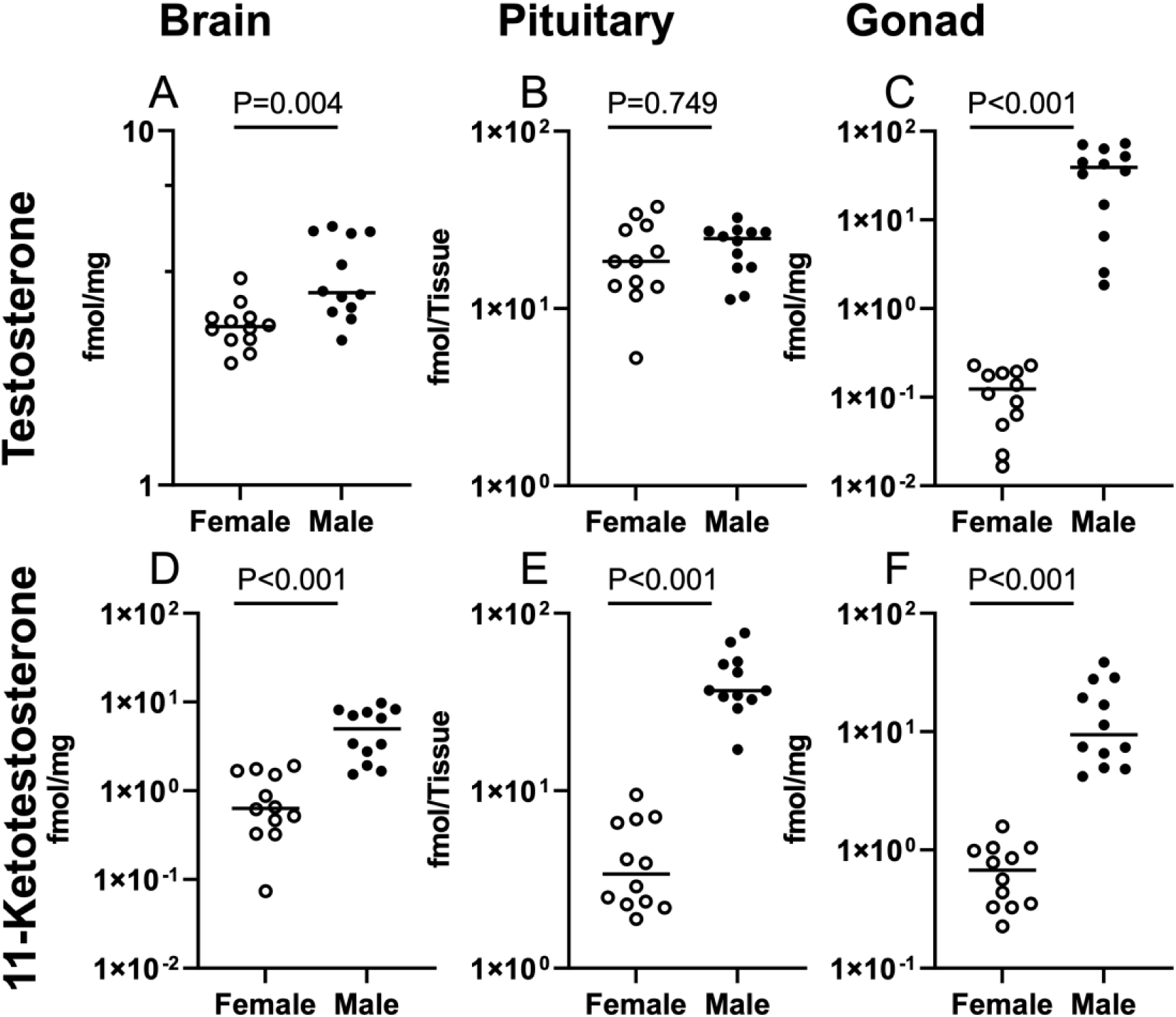
Differences in androgen levels between female and male zebrafish. Testosterone (T) in the brain (A), pituitary (B), and gonads (C). 11-keto-testosterone (11-KT) 11-KT in the brain (D), pituitary (E) and gonads (F). Individual data points for females and males are plotted (n=12) and medians are indicated by the black lines. Note the different scales on the Y-axes. Mann-Whitney U tests were performed. The *p*-values are presented on each graph. See main text for other details.

We calculated the T/11-KT (Table 2) ratio to indicate androgenic conversion steps (steroid figure). Females exhibited 4.32-fold higher T/11-KT ratios in the brain (*z* =3.047, *p*=0.002) and 9.92-fold higher in the pituitary (*z* =4.128, *p*<0.001). In the testes, the T/11-KT ratio was 24-fold higher than in the ovaries (*z*=-3.435, *p*<0.001).

**Table 2.** Steroid conversion indicators in brain, pituitary and gonads of adult zebrafish. Steroid levels were used to calculate the product/parental compound ratios indicated. Results are presented as medians (n=12). Testosterone (T), 11-keto-testosterone (11-KT). Estrone (E1), Estradiol (E2), estriol (E3). Data were analysed using the Mann-Witney U test, and the Z-scores and P values are shown. The female (♀) and male (♂) symbols indicate the sex-bias. See main text for other details.

| RATIOS<br>(median values) |  | ♀ | ♂ | z-scores<br>and p<br>values | Sex<br>Bias |
| --- | --- | --- | --- | --- | --- |
| Brain | T/11-KT | 4.348 | 1.005 | $z=3.047$,<br>$p=0.002$ | ♀ |
| | T/E2 | 0.072 | 0.370 | $z=3.435$,<br>$p<0.001$ | ♂ |
| | E1/E2 | 2.404 | 6.024 | $z=-2.569$,<br>$p=0.011$ | ♂ |
| | E2/E3 | 0.278 | 0.095 | $z=2.973$,<br>$p=0.003$ | ♀ |
| Pituitary | T/11-KT | 5.453 | 0.550 | $z=4.128$,<br>$p<0.001$ | ♀ |
| | T/E2 | 0.086 | 0.179 | $z=-1.241$,<br>$p=0.215$ | none |
| | E1/E2 | 1.253 | 0.395 | $z=2.569$,<br>$p=0.011$ | ♀ |
| | E2/E3 | 0.170 | 0.131 | $z=0.087$,<br>$p=0.928$ | none |
| Gonad | T/11-KT | 0.131 | 3.147 | $z=-3.435$,<br>$p<0.001$ | ♂ |
| | T/E2 | 0.011 | 7.703 | $z=-4.070$,<br>$p<0.001$ | ♂ |
| | E1/E2 | 0.465 | 1.367 | $z=-2.742$,<br>$p=0.006$ | ♂ |
| | E2/E3 | 0.962 | 0.184 | $z=2.742$,<br>$p=0.006$ | ♀ |

Sex differences in the estrogens were apparent in all tissues (Fig. 3). Estrone levels were somewhat higher in the brains of females, being 1.74 times that in males, however, this was not statistically different (*z*=1.472, *p*=0.142). In females, pituitary E1 levels were 3.50 times higher (*z*=2.973, *p*<0.003) than in males. In contrast, testes E1 levels were 4.02 times higher than in ovaries (*z*=2.677, *p*<0.009). E2 levels were higher in females than in males for all tissues. However, only the brain exhibited a clear 3.76-fold difference (*z*=3.262, *p*=0.001). E2 levels in female pituitaries and ovaries were respectively 1.56-(*z*=1.126, *p*=0.259) and 2.15-fold *z*=1.29904, *p*=0.194) than in male tissues, these differences were not statistically different.

**Fig. 3.**
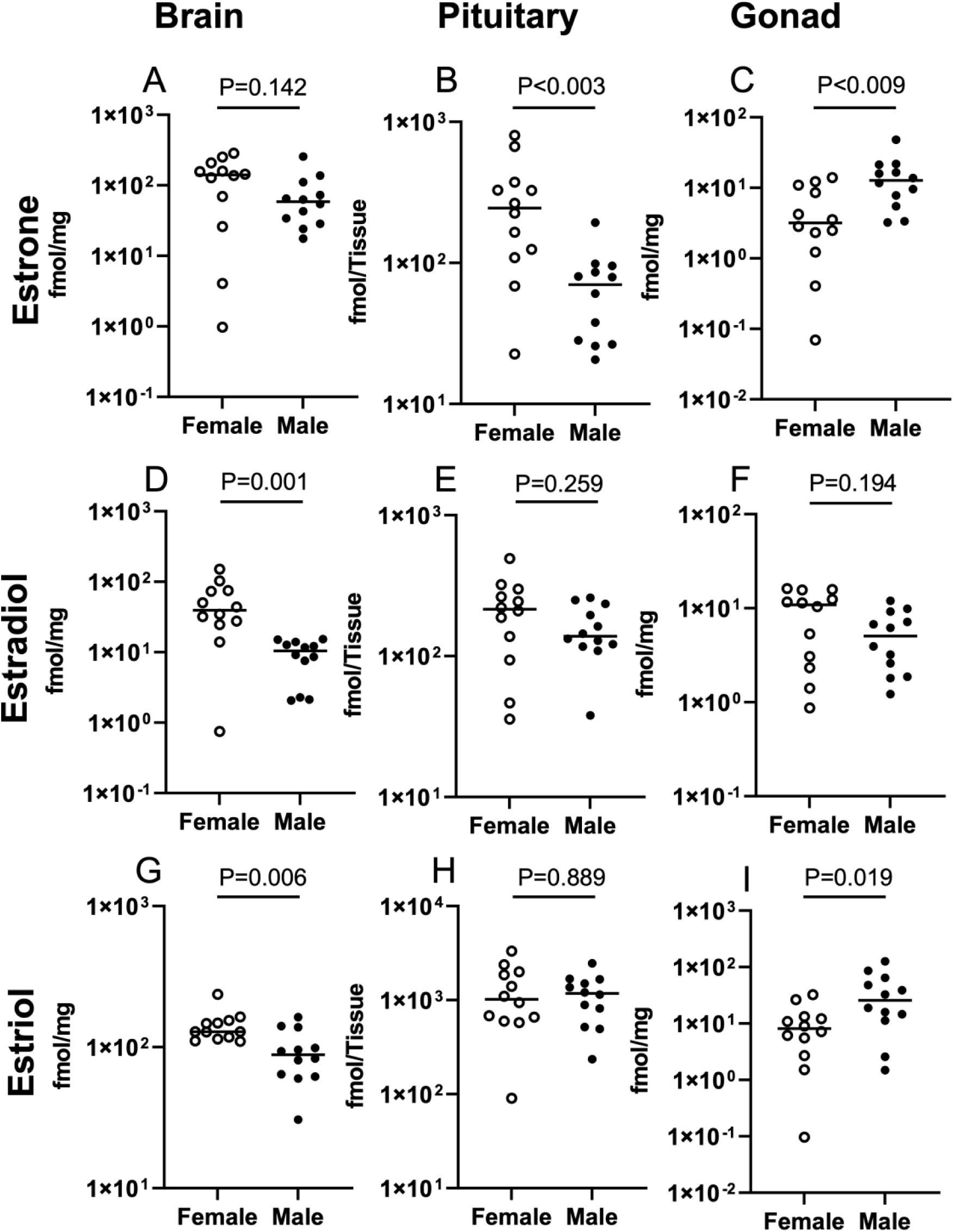
Differences in the levels of estrogens in female and male zebrafish. Estrone (E1) in the brain (A), pituitary (B), and gonads (C). Estradiol (E2) in the brain (D), pituitary (E), and gonads (F). Estriol (E3) in the brain (G), pituitary (H), and gonads (I). Individual data points for females and males are plotted (n=12). Individual data points for females and males are plotted (n=12) and medians are indicated by the black lines. Note the different scales on the Y-axes. Mann-Whitney U tests were performed. The *p*-values are presented on each graph. See main text for other details.

The major estrogenic metabolite E3 was also different between females and males. The levels of E3 in female brains were 1.46 times higher (*z*=2.742, *p*=0.006) than in male brains. There was no difference in E3 levels in the pituitaries (*z*=0.144, *p*=0.889). In contrast, the levels of E3 in the testes were 3.18 times higher (*z*=22.338, *p*=0.019) than in the ovaries.

We calculated several ratios to gain insight into potential sex differences in the metabolism of the estrogens (Table 2). The E1/E2 ratio was 2.51-fold higher in male brains (*z*=-2.569, *p*=0.011), and 2.94 times higher in testes versus ovaries (*z*=-2.742, *p*=0.006). In contrast, the pituitary E1/E2 ratio was 3.17 times higher (*z*=2.569, *p*=0.011) in females compared to males. The E2/E3 ratios were generally higher in females than males. Whereas the E2/E3 ratios were respectively 2.93 (*z*=2.973, *p*=0.003) and 5.22 (*z*=2.742, *p*=0.006) times higher in female brains and ovaries, it was not different in the pituitaries (*z*=0.087, *p*=0.928).

We calculated the T/E2 ratios to indicate T conversion to E2 via the aromatase enzymes (Fig. 1). Specifically, the T/E2 ratio was 680.1 times higher in testes (*z*=-4.070, *p*<0.001) than in ovaries. A less dramatic yet significant 5.13-fold difference (*z*=3.435, *p*<0.001) in the T/E2 ratio was also evident in the male brain. On the other hand, the T/E2 ratio in male pituitaries was not different (*z*=-1.241, *p*=0.215) from that in females.

### Sex differences in the levels of six classical reproductive neuropeptides

We quantified sex differences in the well-known neuropeptides involved in reproduction, Levels of Gnrh2 and Gnrh3 are presented in Fig. 4, There was no significant sex difference (*z*=0.202, *p*=0.842) in the median Gnrh2 levels in the brain, while Gnrh2 levels were 1.91-fold higher in female pituitary (*z*=2.246, *p*=0.024) and 17.83-fold higher in testes than ovaries (*z*=-4.128, *p*<0.001). In the female brain, median Gnrh3 levels were 4.65-fold higher than that in the male (*z*=4.070, *p*<0.001). There were no sex differences (*z*=-0.144, *p*=0.889) evident in pituitary Gnrh3 levels. Testes Gnrh3 levels were 7.61-fold higher than in the ovaries (*z*=-3.376, *p*<0.01).

**Fig 4.**
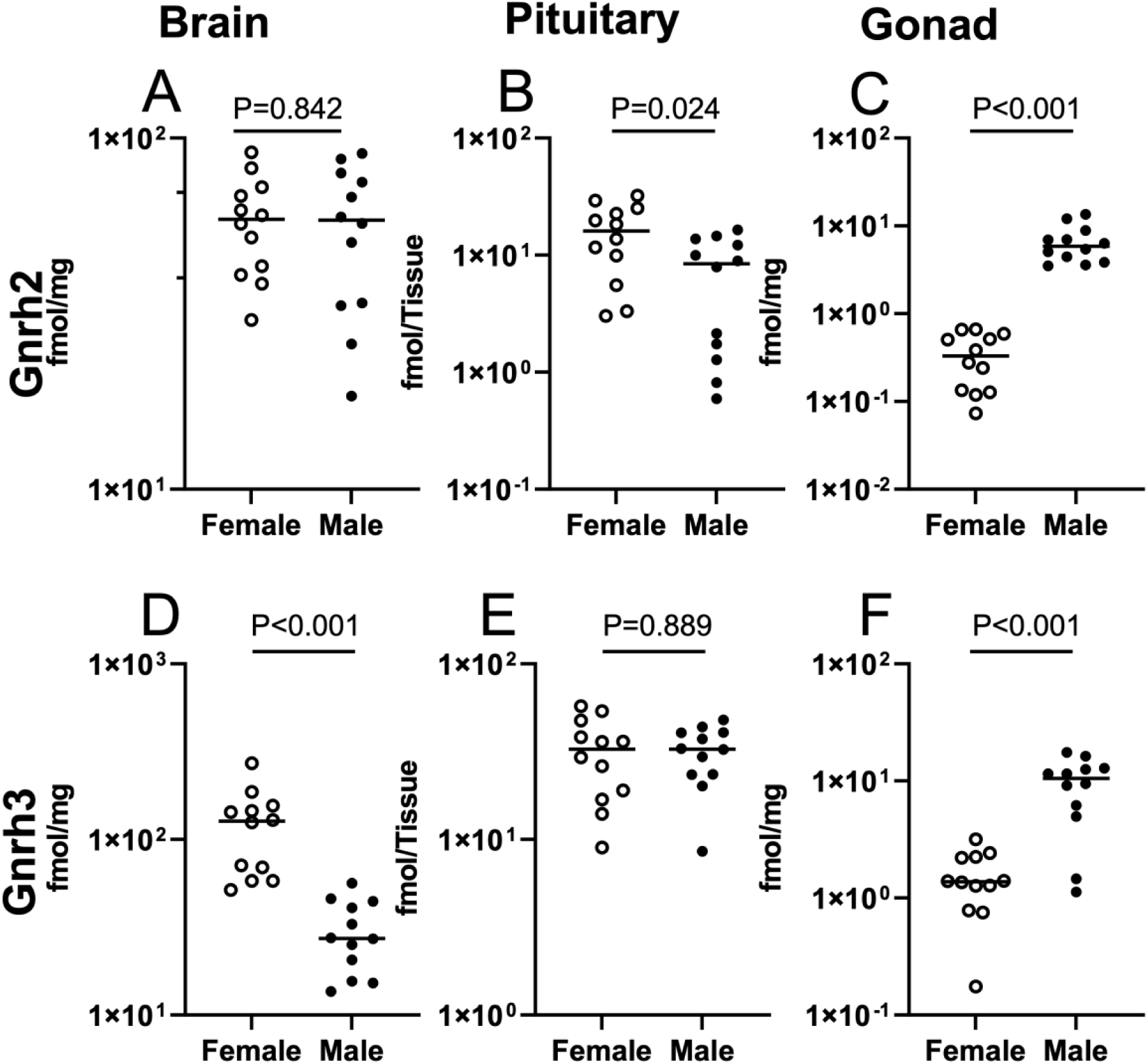
Differences in the levels of gonadotropin-releasing hormone 2 (Gnrh2) and Gnrh3 in female and male zebrafish. Gnrh2 in the brain (A), pituitary (B), and gonads (C). Gnrh3 in the brain (D), pituitary (E), and gonads (F). Individual data points for females and males are plotted (n=12) and medians are indicated by the black lines. Note the different scales on the Y-axes. Mann-Whitney U tests were performed. The *p*-values are presented on each graph. See main text for other details.

Oxytocin and Avp, also respectively known as isotocin and vasotocin in teleosts (Table 1), were quantified (Fig. 5). Oxytocin levels in the brain and pituitary were respectively 2.39-fold (*z*=2.858, *p*=0.004) and 2.18-fold (*z*=2.281, p=0.023) higher in females than that in males. In contrast, Oxt levels were 12.8-fold higher in testes (*z*=-4.128, *p*<0.001) than in ovaries. In a similar pattern to Oxt, Avp levels in the brain and pituitary were respectively 1.33-fold (*z*=3.031, *p*=0.002) and 1.9-fold (*z*=3.551, *p*<0.001) higher in females than that in males (p<0.05). Additionally, Avp levels were 18.74-fold higher (*z*=-4.128, *p*<0.001) in testes compared to ovaries.

**Fig 5.**
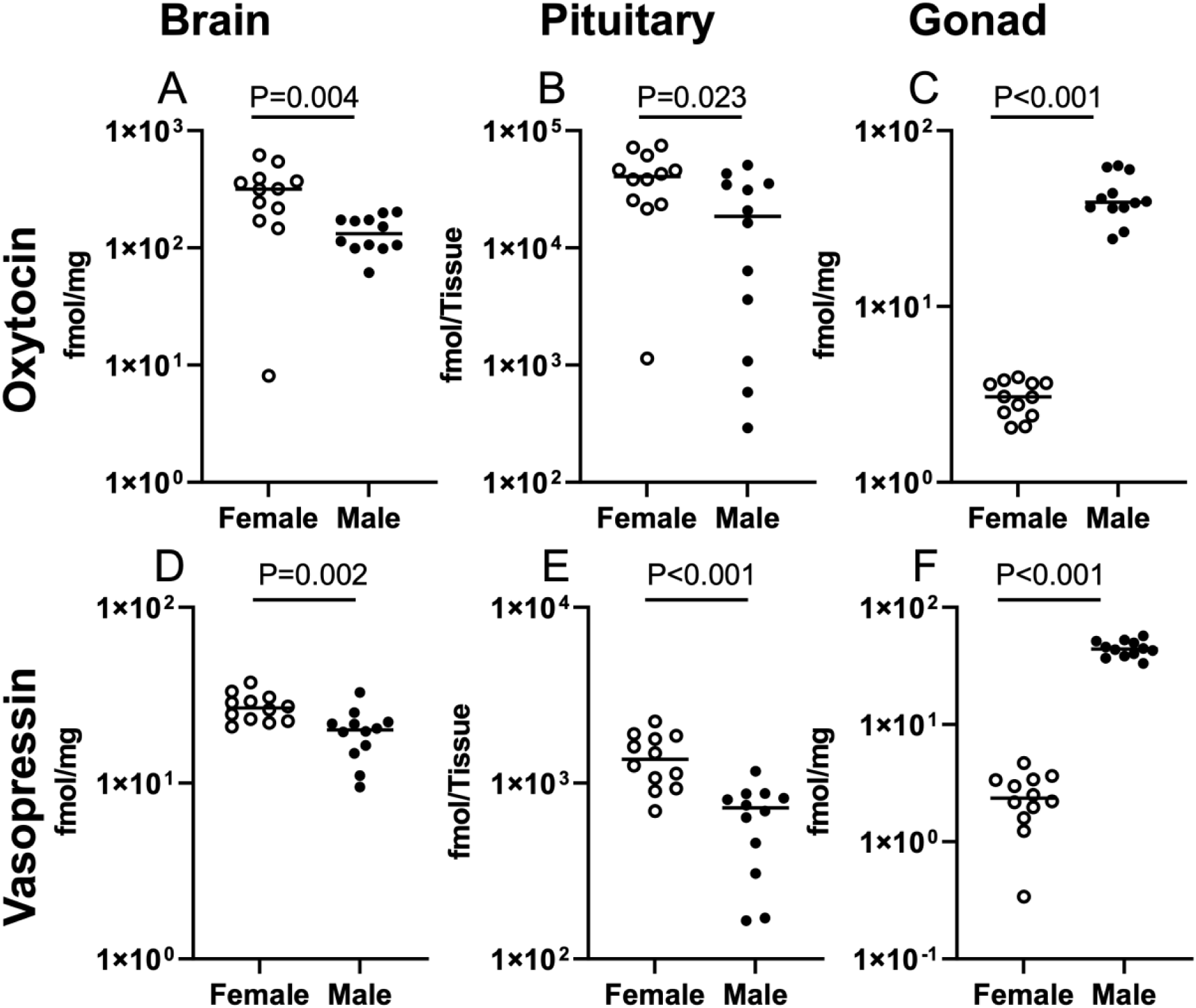
Differences in oxytocin (Oxt) and vasopressin (Avp) levels in female and male zebrafish. Oxt in the brain (A), pituitary (B), and gonads (C). Avp brain (D), pituitary (E) and gonads (F). Individual data points for females and males are plotted (n=12) and medians are indicated by the black lines. Note the different scales on the Y-axes. Mann-Whitney U tests were performed. The *p*-values are presented on each graph. See main text for other details.

Levels of Kiss1 and Kiss2 are presented in Fig. 6. There were no sex differences in the median levels of Kiss1 (*z*=-1.93412, *p*=.054) and Kiss2 (*z*=1.645, *p*=0.099) in the brains. Similarly, we found no evidence for sex differences in Kiss1 (*z*=0.779, *p*=0.435) and Kiss2 (*z*=1.812, *p*=0.069) in pituitaries. In contrast, median values of both Kiss1 and Kiss2 were respectively 12.87 (*z*=-3.89711, *p*<0.001) and 13.48 (*z*=-4.129, *p*<0.001) times higher in testes than ovaries.

**Fig 6.**
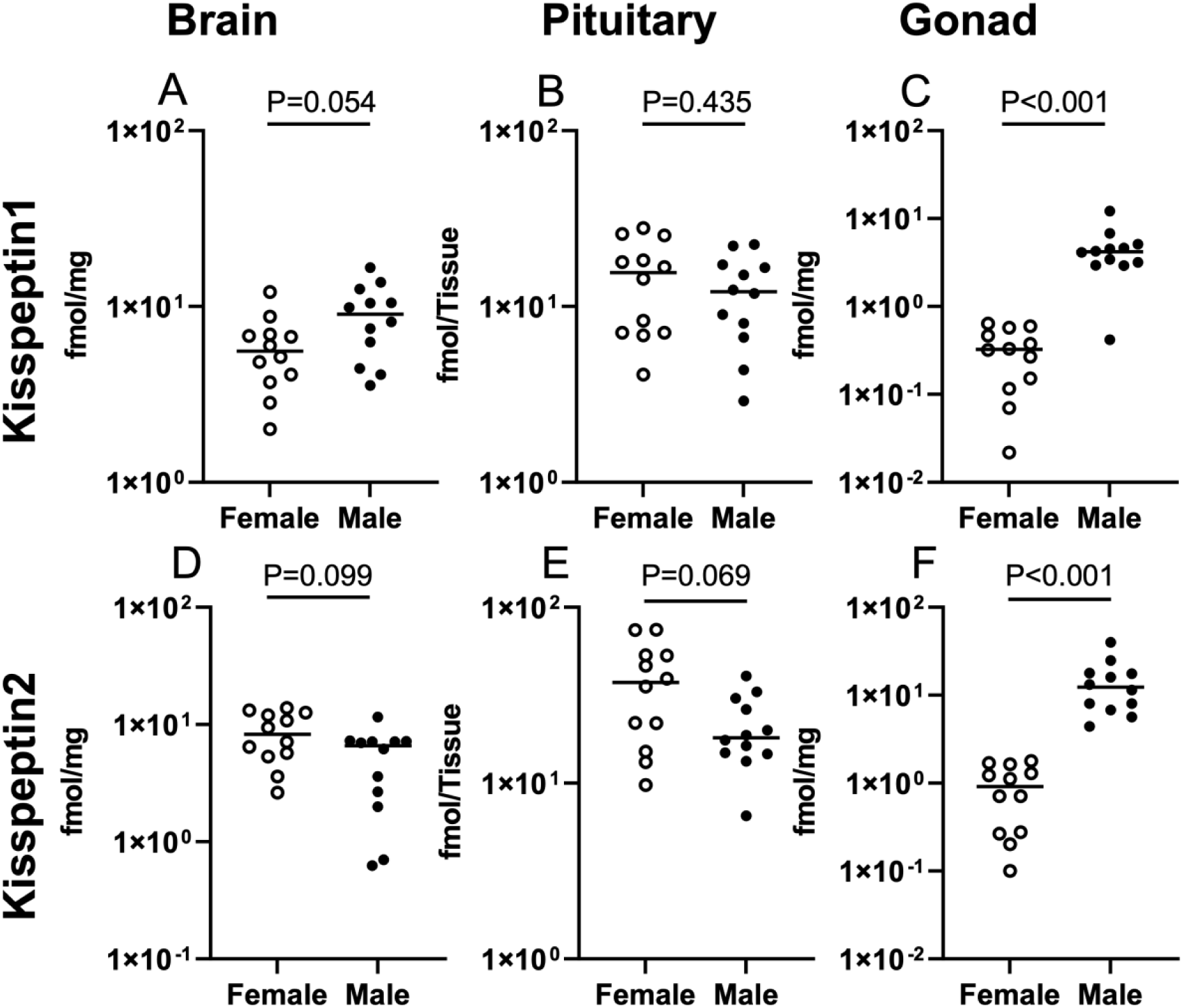
Differences in kisspeptin1 and kisspeptin2 levels in female and male zebrafish. Kiss1 in the brain (A), pituitary (B) and gonads (C). Kiss2 in the brain (D), pituitary (E) and gonads (F). Individual data points for females and males are plotted (n=12) and medians are indicated by the black lines. Note the different scales on the Y-axes. Mann-Whitney U tests were performed. The *p*-values are presented on each graph. See main text for other details.

### Sex differences in SNa1-34, SNb1-31 and related fragmental peptides

SNa1-34 was detected in all 3 tissues in both females and males (Fig 7). The levels of SNa1-34 female brains and pituitaries were respectively 4.15 (*z*=4.128, *p*<0.001) and 1.43 (*z*=3.089, *p*=0.002) times higher than in males. In contrast, SNa1-34 was 13.73 times higher (*z*=-4.128-, *p*<0.001) in the testes than ovaries.

**Fig 7.**
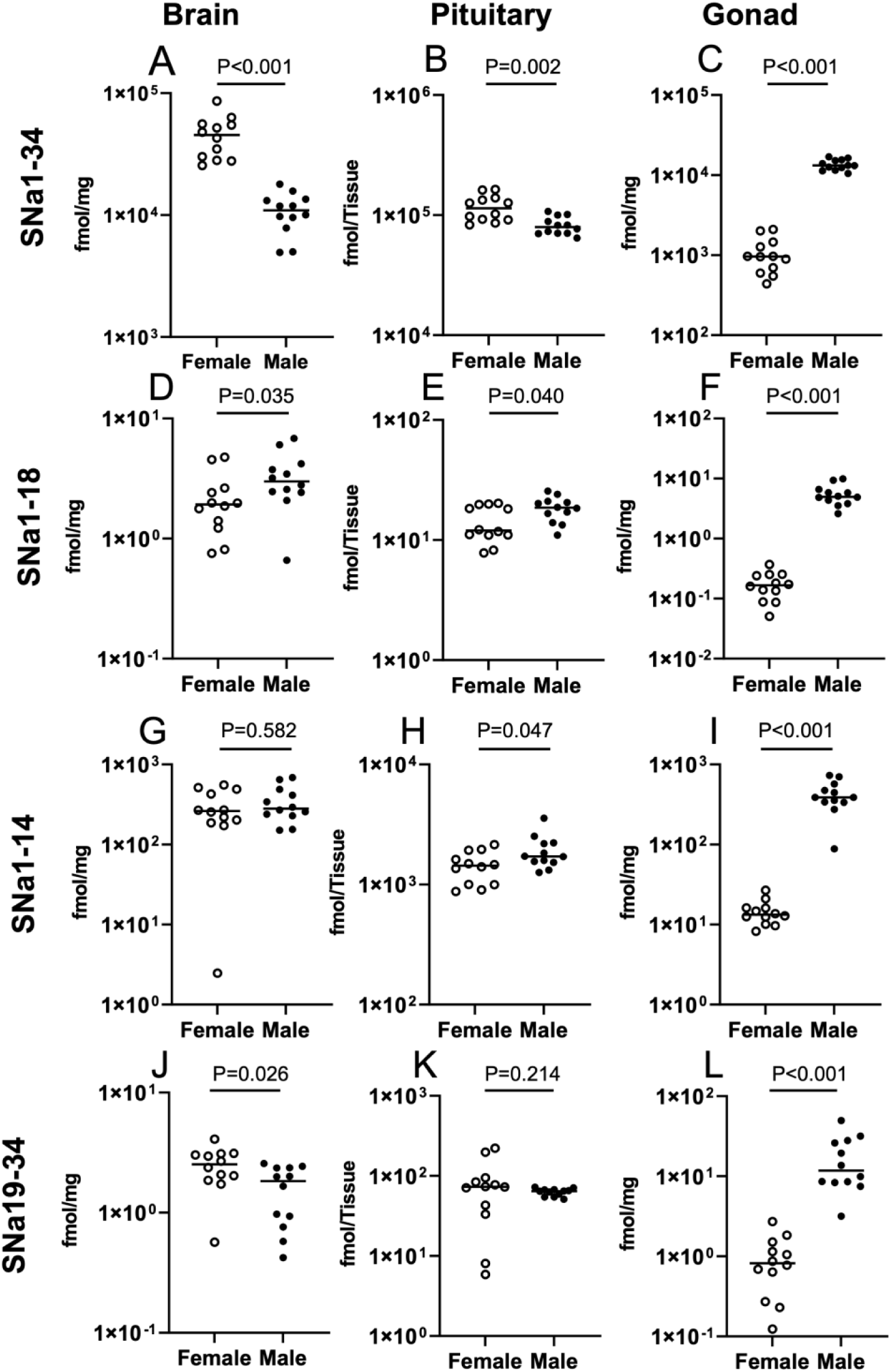
Differences in secretoneurin A (SNa1-34) and fragmental SNa peptide levels in female and male zebrafish. SNa1-34 in the brain (A), pituitary (B) and gonads (C). SNa-18 in the brain (D), pituitary (E) and gonads (F). SNa1-14 in the brain (G), pituitary (H) and gonads (I). SNa19-34 in the brain (J), pituitary (K) and gonads (L). Individual data points for females and males are plotted (n=12) and medians are indicated by the black lines. Note the different scales on the Y-axes. Mann-Whitney U tests were performed. The *p*-values are presented on each graph. See main text for other details.

SNa1-34 can be processed into numerous smaller fragmental peptides. We quantified the three most abundant. Firstly, the N-terminal fragment SNa1-18 was lowly abundant, representing on average ∼0.02% of the levels of SNa1-34 over all tissues. Nevertheless, SNa1-18 was higher in males than females. Levels of SNa1-18 were respectively 1.56 (*z*=-2.107, *p*=0.035), 1.55 (*z*=-2.049, *p*=0.040) and 29.77 (*z*=-4.128, *p*<0.001) times higher in male brains, pituitaries and testes. Secondly, the SNa1-14 fragment was the most abundant, representing on average ∼2.22% of the levels of SNa1-34 over all tissues. Levels of SNa1-14 were similar in female and male brains (*z*=-0.548, *p*=0.582). The SNa1-14 peptide was 1.19 times higher in male versus female pituitaries (*z*=-1.992, *p*=0.047). In testes, SNa1-14 was 29.25 times higher (*z*=-4.128, *p*<0.001) than in ovaries. Thirdly, the C-terminal SN19-34 fragment was also lowly abundant, representing on average ∼0.07% of the levels of SNa1-34 over all tissues. The SNa19-34 fragment was respectively 3.41 (*z*=-2.223, *p*=0.026) and 14.35 (*z*=-4.128, *p*<0.001) times higher in the brains and gonads of males compared to females. In contrast, there were no sex differences evident for SNa19-34 in pituitaries (*z*=-1,241, *p*=0.214).

Next, we calculated the ratios of fragmental SNa peptide levels to indicate potential processing differences. Most SNa fragmental ratios were significantly higher in males than females (see Table 3 for statistical results). For the brain, the relative production of SNa1-18, SNa19-34 and SNa1-14 were 7.52, 3.41 and 4.16-fold higher in males (Table 3). In the pituitary, the relative production of SNa1-18 and SNa1-14 were 2.38 and 1.66 times higher in males (table). Similarly in the testes, the relative production of SNa1-18 and SNa1-14 were 2.30 and 2.00 times higher than in the ovaries (table).

**Table 3.**
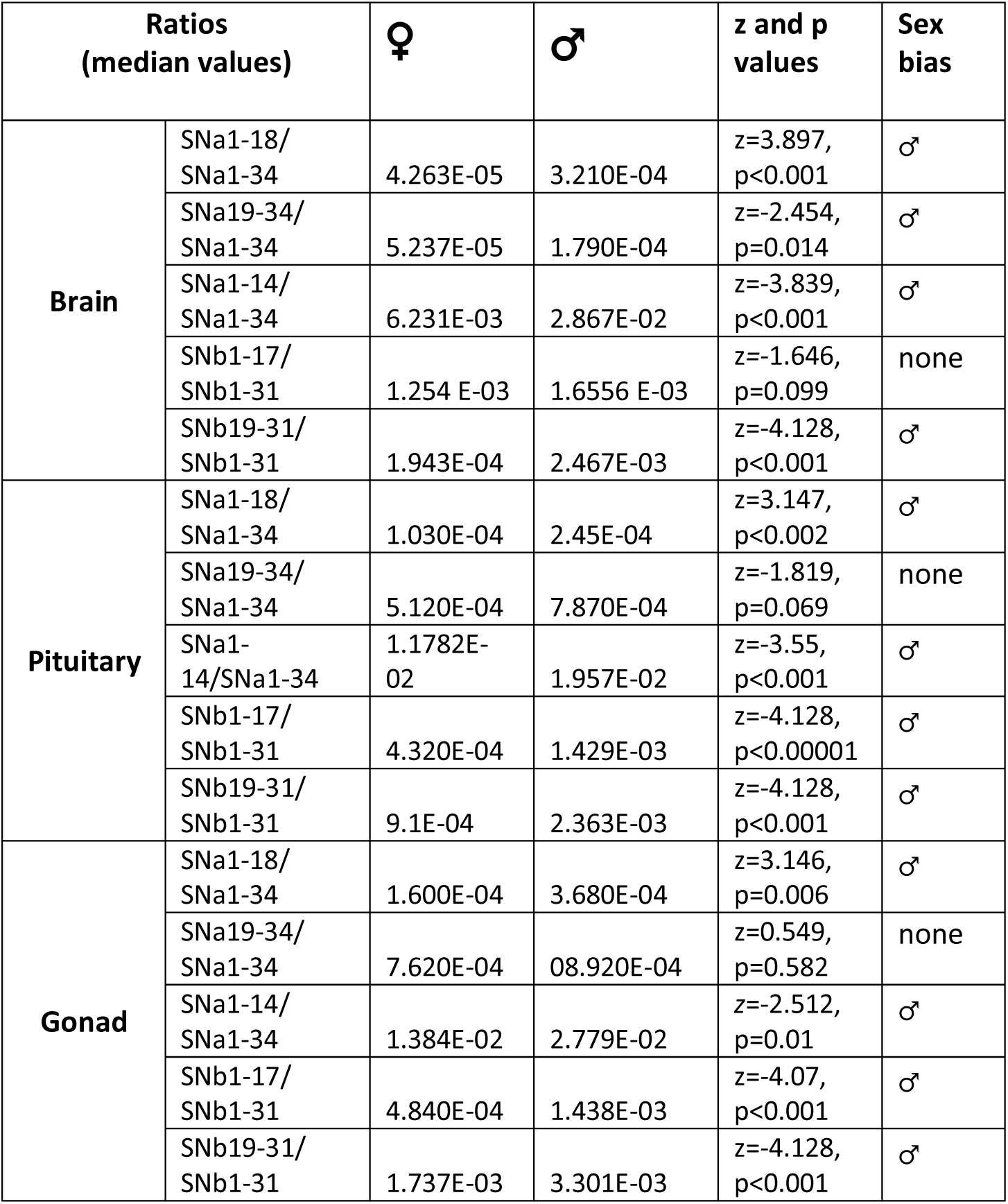
Ratios of SNa and SNb fragmental peptides derived from SNa1-34 and SNb 1-31. Results are presented as medians (n=12). Data were analysed using the Mann-Witney U test, and the Z-scores and P values are indicated. The male (♂) symbol indicates the sex-bias. See main text for other details.

Clear sex differences in the levels of SNb1-31 were apparent in all tissues. The levels of SNb1-31 female brains and pituitaries were respectively 3.98 (z= 4.128, p <0.001) and 4.12 (z= 4.128, p<0.001) times higher than in males. In contrast, SNb1-31 was 6 times higher (z= -4.128, p<0.001) in the testes than ovaries (Fig 8). These differences should be interpreted with caution, as SNb1-31 measurements approached both the lower and upper limits of quantification due to analytical challenges, suggesting that the observed higher concentrations may represent underestimations of true physiological levels.

**Fig 8.**
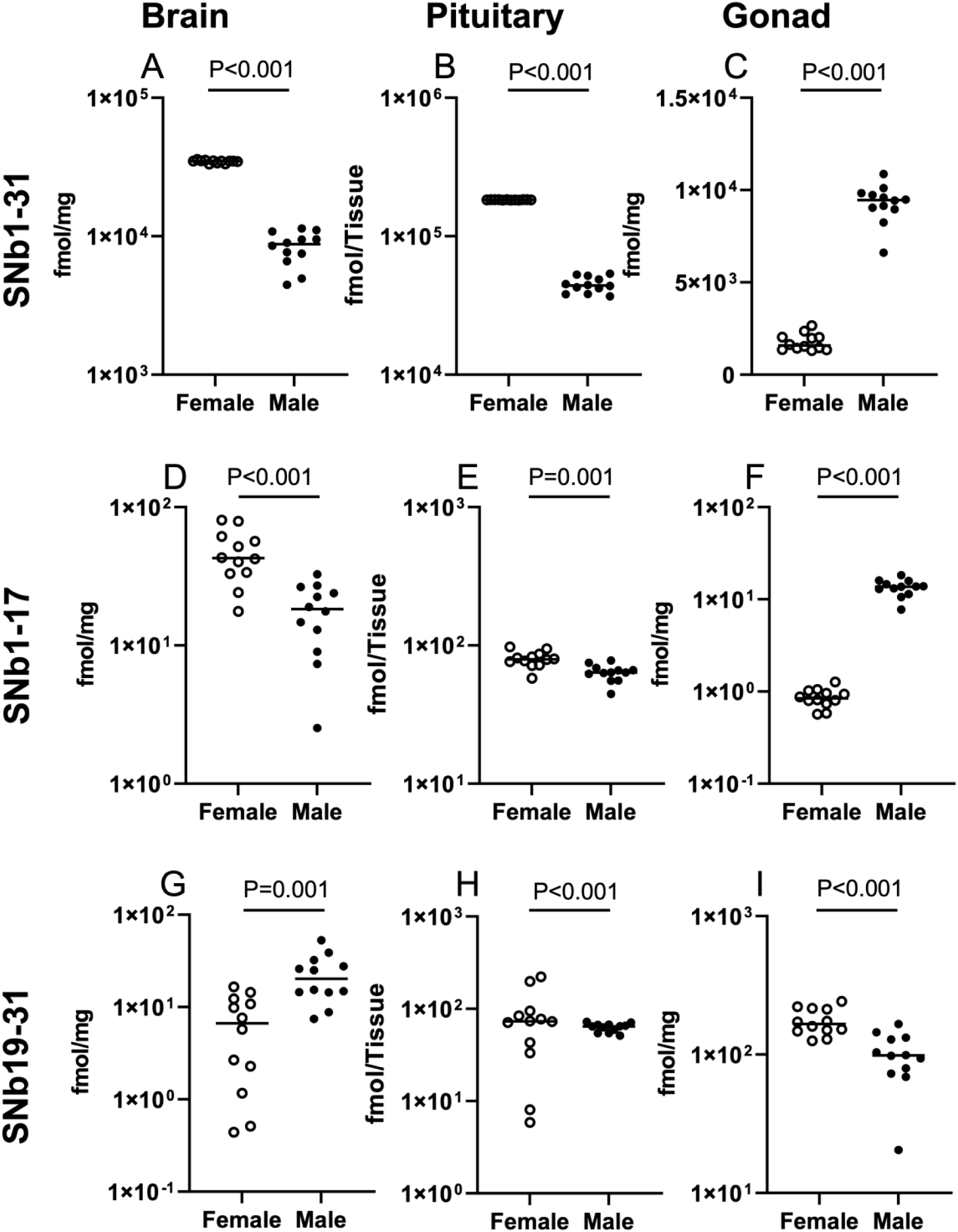
Differences in SNb1-31 and fragmental SNb peptide levels in female and male zebrafish. SNb1-31 in the brain (A), pituitary (B) and gonads (C). SNB1-17 brain (D), pituitary (E) and gonads (F). SNb19-31 in the brain (G), pituitary (H) and gonads (I). Individual data points for females and males are plotted (n=12) and medians are indicated by the black lines. Note the different scales on the Y-axes. Mann-Whitney U tests were performed. The *p*-values are presented on each graph. See main text for other details.

As with SNa1-34, SNb1-31 can be further processed into numerous smaller fragmental peptides, and we quantified the two most abundant. Firstly, the N-terminal fragment SNb1-17 was lowly abundant, representing on average ∼0.12% of the levels of SNb1-31 over all tissues. While this is very low, SNb1-17 exhibited a similar pattern of sex difference as its parent peptide.

The levels of SNb1-17 female brains and pituitaries were respectively 2.33 (z=3.608, p<0.001) and 1.25 (z=3.262, p=0.001) times higher than in males. In contrast, SNb1-31 was 16.29 times higher (z= -4.128, p<0.001) in the testes than ovaries (Fig 8). Secondly, the C-terminal fragment SNb19-31 was lowly abundant, representing on average ∼0.2% of the levels of SNb1-31 over all tissues. The levels of SNb19-31 male brains and gonads were respectively 3.03 (z=-3.204, p=0.001) and 9.90 (z= -4.128, p<0.001) times higher than in males. In contrast, SNb19-31 was 1.69 times higher (z= 3.493, p<0.001) in female pituitaries (Fig 8).

Most SNb fragmental ratios were also significantly higher in males than females (see Table 3 for statistical results). While the relative production of SNb1-17 was not different between the sexes, the SNb19-31/SNb1-31 ratio was 12.7 times higher in males than females (Table 3). In the pituitary, the relative production of SNb1-17 and SNb19-31 were 3.31 and 2.60 times higher in males. In the testes, the relative production of SNb1-17 and SNb19-31 were respectively 2.97 and 1.90 times higher than in ovaries.

In brain tissue (Fig 9A, B), PC1 and PC2 collectively explained 48.49% (36.10% and 12.39%) and 49.80% (36.89% and 12.91%) of total variance for analyses excluding and including SNa/SNb peptides, respectively. Similar analyses in pituitary (Fig 9C, D) explained 49.39% (27.40% and 21.99%) and 44.85% (29.14% and 15.71%) of variance, and in gonads (E, F) explained 67.24% (55.23% and 12.01%) and 73.00% (65.10% and 7.90%) of variance. In the first case when the SNs were excluded, there was reasonable separation by sex for brain and gonads. For pituitary the sexes were less distinct, although for female tissues there was a wider distribution along both axes (e.g., PC1 vs. PC2). Markedly, inclusion of SNa1-34, SNb1-31 and related fragmental peptides amplifies sex differences in all tissues. For brain and pituitary, inclusion of SNs improves separation (PC1) of females from males, with PC2 highlight individual variation. For the gonads, females exhibited far less variability compared to males.

**Figure 9.**
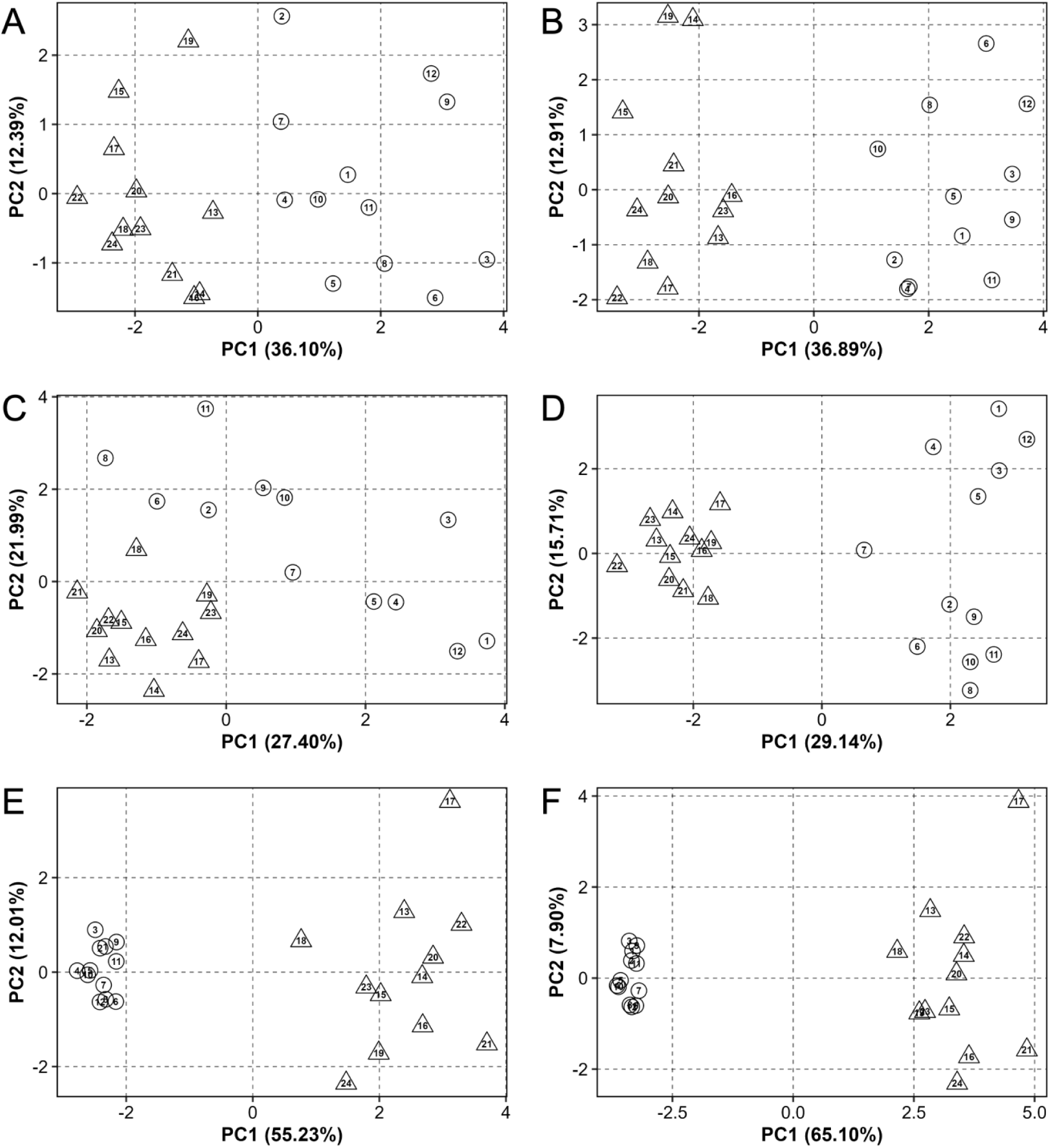
Principal component analysis (PCA) of steroid and neuropeptide and hormone levels in brain (A, B), pituitary (C, D), and gonads (E, F) of male and female zebrafish. Panels A, C, and E represent PCA results excluding SNa, SNb and their fragments, while panels B, D, and F show analysis including all quantified peptides. Numbers represent individual samples, with circles indicating females and triangles indicating males. The spatial distribution of samples reflects their relative similarities in peptide and hormone expression patterns, with greater distances indicating larger differences.

## Discussion

Few studies in fish have documented sex differences in classical sex steroids and neuropeptides. Establishing this was necessary before examining the SNs—an emerging reproductive hormone family derived from proteolytic processing of Scg2 (Mitchell, Mikwar et al., 2020; Fleming et al., 2024; Peng et al., 2025). This was made possible by our ability to simultaneously detect and quantify lipophilic steroids and hydrophilic peptides in the same small tissues using novel extraction and LC-MS methodologies.

There were clear male-dominant sex differences in testicular androgens. Levels of T and 11-KT were approximately 300- and 10-fold higher in testes than ovaries, respectively. Testosterone was also somewhat higher in male brains, while pituitary T levels were similar between sexes. In contrast, brain and pituitary 11-KT levels showed a more striking male bias, being ∼8 and ∼10 times higher than in females. In teleosts, 11-KT is the primary bioactive androgen, synthesized from T via Cyp11c1 and Hsd11b2 (Fig. 1). Ratios of T/11-KT were used to assess tissue and sex differences in androgenic conversion. Lower T/11-KT ratios in male brain and pituitary suggest greater conversion of T to 11-KT compared to females. The higher testicular T/11-KT ratio relative to ovaries reflects the predominance of T, due to its very high levels in testes. Regarding reproductive functions, 11-KT promotes spermatogenesis in Japanese eel (Miura et al., 1991) and sexual development in zebrafish (Zhang et al., 2020). In ovaries, 11-KT induces primary follicle development in coho salmon (Monson et al., 2017) but inhibits follicular progression in zebrafish by blocking the granulosa-theca cell cycles (Sousa et al., 2016). These findings support our ovarian 11-KT measurements and suggest roles in ovarian and follicular development. In tilapia, 11-KT also induces GnRH3 neuron proliferation in the brain (Narita et al., 2018), possibly reflecting sex-specific androgen-to-estrogen ratios.

The distinct male-biased androgen levels contrasted with estrogens. While E1, E2, and E3 were present across tissues, brain levels of E2 and E3 were higher in females. The E1/E2 ratio suggests E1 predominates in male brains, whereas E2 is dominant in females. E2/E3 ratios, lower in males, indicate more active E2-to-E3 conversion in females. The T/E2 ratio was lower in female brains, reflecting more active androgen-to-estrogen conversion. Zebrafish brains show high de novo steroidogenesis, converting pregnenolone to both androgens and estrogens (Diotel et al., 2018). Expression of *cyp19a1b* and aromatase activity are generally higher in female teleost brains. High estrogen and aromatase levels participate in a local positive feedback loop in radial glial cells—the sole site of brain aromatase B (Diotel et al., 2018). In a glial cell assay, E2 more potently induces aromatase B via all three nuclear estrogen receptors (Le Page et al., 2005). Estradiol also inhibits neurogenesis in zebrafish, which shows sex- and brain region-specific variation (Diotel et al., 2018). Inhibiting aromatase disrupts female but not male social approach behavior (Lindley et al., 2025). *Cyp19a1b-/-* females take longer to oviposit than WT, while male *cyp19a1a-/-* and *cyp19a1b-/-* mutants show no difference from WT males (Shaw, Lu et al., 2023; Shaw, Therrien et al., 2023). Interestingly, *esr2b-/-* females in medaka retain ovarian function but reject males and court females; *esr2b-/-* males spawn normally (Nishiike et al., 2021). Differences in brain steroids are therefore linked to sex-dependent regulation of fundamental processes in the adult brain.

Pituitary cells may be regulated by locally produced sex steroids in addition to classical gonadal feedback. The most striking pituitary steroid differences were a high T/11-KT ratio in males and a moderately higher E1/E2 ratio in females, indicating active steroid conversion in both sexes. Isolated gonadotropic cells from African catfish (*Clarias gariepinus*) can convert androstenedione into E1 (de Leeuw et al., 1985). Species-specific expression of multiple ERs, ARs, and aromatase suggests complex regulatory mechanisms for gonadotrophs and somatotrophs (Callard et al., 1988; Fontaine et al., 2020). In medaka, E2 in females and 11-KT in males reduced Fsh cell number after gonadectomy-induced increases. Follicle stimulating hormone cell hyperplasia followed gonadectomy, with new Fsh cells arising from mitosis and transdifferentiation of Tsh cells in females—but not males (Royan et al., 2023). These authors did not distinguish estrogen versus androgen contributions. In goldfish, T (via aromatization to E2) enhanced pituitary Lh release in response to exogenous Gnrh agonist in both sexes (Trudeau et al., 1991). In vitro, T—but not 11-KT—increased pituitary Gnrh responsiveness (Trudeau, Murthy et al., 1993). Testosterone also enhanced GABA- and taurine-stimulated Lh release in vivo (Trudeau et al., 1993) and induced gonadotropin production (Habibi & Huggard, 1998). E2 and *esr* gene expression (*esr1*, *esrra*, *esr2b*) in LH gonadotrophs show estrogens act directly on the pituitary to induce Lhb synthesis in female zebrafish (Li et al., 2018; Tanaka et al., 2024).

Our data on sex differences in steroids laid the foundation for a detailed analysis of reproductive peptides. Zebrafish and many cyprinids express both Gnrh2 and Gnrh3, which we measured in brain, pituitary, and gonads. As both Gnrh2 and Gnrh3 neurons innervate the pituitary, their detection reflects peptide levels in nerve terminals at the release site (Trudeau & Somoza, 2020). Female-biased pituitary Gnrh2 and brain Gnrh3 levels were observed, while testes had higher levels of both than ovaries. Sex differences in Gnrh function have been reported in several species. Gnrh2 and Gnrh3 stimulate Lh production and secretion in many teleosts (Zohar et al., 2021), with similar potencies via the type 2 Gnrh receptor (Tello et al., 2008), the main form in zebrafish Lh gonadotrophs (Tanaka et al., 2024). Intracebroventricular injection studies in goldfish revealed important stimulatory roles for the Gnrhs in female spawning behavior; however, having no apparent effects on male behaviour (Volkoff & Peter, 1999). Female zebrafish with Gnrh2 mutations had smaller oocytes and produced embryos with higher mortality than WT (Marvel et al., 2019). Fasting zebrafish showed a 4-fold increase in Gnrh2 fiber length to the pituitary and a 6-fold decrease in Gnrh3 fibers (Marvel et al., 2021). Breeding success after 14-day fasting was significantly higher in WT than Gnrh2 mutants of both sexes, supporting that Gnrh2 modulates reproduction via a different pathway than Gnrh3.

The most obvious differences were the relatively higher levels of Gnrh2 and Gnrh3 in the testes compared to ovaries, yet intragonadal Gnrh peptides play important roles in both sexes. In goldfish, both Gnrh2 and Gnrh3 directly stimulate germinal vesicle breakdown (GVBD) but inhibit hCG-induced GVBD (Pati & Habibi, 2000). These peptides alone do not affect follicular cell T production but reduce Lh-stimulated T. Gnrh3 is expressed in follicular cells and stimulates GVBD as well as caspase-3 activity in late-vitellogenic follicles to induce oocyte maturation (Fallah & Habibi, 2020). In vitro studies in zebrafish showed Gnrh3 stimulates testicular cell development (Fallah et al., 2020). Gnrhs are expressed in different spermatogonia and Leydig cells; both Gnrh2 and Gnrh3 stimulate spermatogenesis and T production, with Gnrh2 being more potent. Gnrh3 also contributes to sex differentiation by regulating PGC proliferation, and its knockout promotes a male-biased sex ratio (Feng et al., 2020). Remarkably, mutation of Gnrh2 and Gnrh3 does not affect overall reproductive performance in adult females or males, suggesting other neuropeptides may play a critical role in zebrafish (Whitlock et al., 2019).

Levels of Oxt and Avp were higher in female brain and pituitary, while both were much more abundant in zebrafish testes than ovaries. In the catfish *Heteropneustes fossilis*, brain Avp shows season-dependent sex differences: female-biased pre-spawning, then male-biased at and after spawning. Unlike zebrafish, catfish testes have lower Avp than ovaries (Singh & Joy, 2008). In medaka, the most prominent sex difference was in *avt* expression in the nucleus posterior tuberis (NPT) and posterior nucleus ventral tuberis (NVT) of the hypothalamus, where expression is male-specific (Kawabata et al., 2012). Male-biased *oxt* expression in the parvocellular preoptic nucleus is driven by gonadal androgens (Yamashita et al., 2017), as is male-specific *avp* expression in the tuberal hypothalamus (Kawabata-Sakata et al., 2024). Functionally, there is a distinct sex difference in medaka for Oxt. Using various knockout models, it was determined that females lose mate preference while mutant males gain mate preference for familiar mates (Yokoi et al., 2020). In zebrafish, Avp-deficient couples produce significantly fewer fertilized eggs per clutch than wildtypes (Ramachandran et al., 2023). Crossbreeding experiments show this reproductive phenotype is female-dependent, since Avp-deficient males reproduce normally with wildtype females. Thus, there are clear sex differences in Oxt and Avp production and function in several fish species.

Higher Kiss1 levels in the male brain were evident. In mammals, kisspeptin regulates reproduction by stimulating Gnrh1 release (Sivalingam et al., 2022). In teleosts, gene mutation studies indicate that kisspeptins and their receptors are dispensable (Tang et al., 2015). In medaka, male-dominant *kiss1* expression was found in the nucleus ventral tuberis of the brain (Kanda et al., 2008). In zebrafish, the most striking difference was the high testicular levels of both Kiss1 and Kiss2. Kisspeptins have direct biphasic effects on ovarian steroidogenesis in human granulosa cells (Fabová et al., 2022), and time- and dose-dependent effects on testicular steroidogenesis in humans and frogs. A defined role of kisspeptins is in autocrine/paracrine control of frog spermatogenesis (Meccariello et al., 2020), but much remains to be clarified about sex differences in Kiss1 and Kiss2 function in teleost gonads.

The emerging stimulatory role of SN family peptides prompted untargeted peptidomic analysis (Peng et al., 2025) to assess natural production and abundance of peptides derived from Scg2 and Scg2b precursors. Our results confirm that SNa1-34 and SNb1-31 sequences match predictions from genomic data. In *scg2a/scg2b* zebrafish double mutants with impaired spawning, i.p. injection of SNa1-34 partially rescued reproductive defects, while SNb1-31 did not (Mitchell, Zhang et al., 2020). Although little is known about SNb1-31 bioactivity, SNb injection increases hypothalamic *gnrh3* expression and pituitary *lhb* and *fshb* levels in grouper (Shu, Yang et al., 2018). In addition to intact SNa1-34 and SNb1-31 peptides, we identified 6 naturally occurring SNa fragments and 17 for SNb (Peng et al., 2025). These findings suggest SNa1-34 and SNb1-31 are not the only biologically active Scg2a/2b-derived peptides. We therefore selected SNa1-14, SNa1-18, SNa19-34, SNb1-17, and SNb19-31 to examine sex differences. Across all tissues and both sexes, the SNa1-14 fragment was the most abundant, representing only ∼2% of measured SNa1-34 levels. All other SNa and SNb fragments were very low but quantifiable.

Of these only SNa1-34 and SNa1-14 have been tested for biological activity in zebrafish, and when injected i.p., SNa1-34 but not SNa1-14 rapidly and robustly increases the expression of telencephalon and hypothalamic *gnrh3*, pituitary *lhb* and *cga*, and ovarian *lhcgr* and *npr* which leads to ovulation (Peng et al., 2025). On the other hand, both peptides similarly stimulate telencephalon *gnrh2* and *oxt* (Peng et al., 2025). There were marked female-biased levels of SNa1-34 and SNb1-31 in brain and pituitary. Moreover, testicular levels of both SNa1-34 and SNb1-31 were high, and at the upper limit of quantification. The first report on measurements of SN levels was for various rat brain regions, with no specification of which sex was under study (Marksteiner et al., 1993). Mammalian SN is 33 amino acids long and is the equivalent to the phylogenetically older SNa1-34 form found in all fish. For SNa1-18 levels were generally higher in male tissues than in female. SNa1-14 levels were higher in male pituitary and testes, and for SNa19-34, brain levels were somewhat higher in females yet were higher in testes that ovaries. The Scg2b precursor arose from the teleost-specific whole genome duplication event (Peng et al., 2025), and when processed yields SNb1-31, SNb1-17 and SNb 19-31. For SNb1-17, levels were higher in female brain and pituitary, but for SNb19-17, levels were higher in male brain and lower in male pituitary, both exhibiting a male bias in the testes. Despite being of low abundance, higher ratios of the fragmental peptides imply male-biased processing across all tissues. In mammals, there is a role for both PC1 and PC2 to cut at dibasic cleavage sites to yield SN; relative involvement of either PC depends on the tissue or cell system studied (Mitchell, Mikwar et al., 2020). The proteases responsible for generating the smaller SN fragments are not known. Nevertheless, we report for the first time, sex differences in the SN family of peptides in brain, pituitary and gonads. Within and between line crossing of *scg2a* and *scg2b* frameshift mutants also revealed potential sex differences. For example, double mutant females are poor ovulators when paired with normal males, whereas the double mutant males have normal fertility but have somewhat reduced offspring survival (Mitchell et al., 2020).

To test whether the SN peptides contribute to sex differences, we performed PCA analyses for each tissue both excluding and including SNa/SNb peptides to determine their impact. Inclusion of the SNs improved separation of females from males across all three tissues, indicating that these peptides contribute significantly to sex-specific hormonal signatures. This places the SN peptide system among other steroid and peptide hormones exhibiting clear sex differences. Future research should consider possible sex differences in the role and regulation of the SN peptides along the brain-pituitary-gonadal axis.

## Author contribution statement

P.D. and C. Lu are co-first authors and performed the experiments, data analysis and writing. V.L.T. provided funding, helped design experiments, performed statistical analysis, and wrote the paper. All authors approved the final version of the manuscript.

## Declaration of interest, Funding and Acknowledgements

The authors have no conflicts of interest to declare. This research was funded by the China Scholarship Council (to DP), the Natural Sciences and Engineering Research Council of Canada Discovery Grant (RGPIN-2021-03174 to V.L.T.), the University of Ottawa Research Chair program (to V.L.T.) The authors acknowledge with appreciation the help of E. Ramsay (ACVS) for care of animals, and Dr. Z. Ning for advice on mass spectroscopy. The senior author recognizes that he works and lives on the unceded Algonquin Territory of the Three Fire Confederacy, Anishinaabewaki.

